# Human induced pluripotent stem cell-derived astrocyte and neuron co-culture model for neuroinflammation modeling

**DOI:** 10.1101/2025.04.13.648609

**Authors:** Boning Qiu, Massimiliano Caiazzo

## Abstract

Access to human neural cells is important for faithful *in-vitro* neural modeling as animals have innate species gaps. However, primary human cells are difficult to retrieve whereas immortalized human cell lines do not guarantee complete authenticity. Differentiation of human induced pluripotent stem cells (hiPSCs) has emerged as a novel approach to derive faithful human neural cells. Here we differentiated hiPSCs into functional midbrain dopaminergic (mDA) neurons and astrocytes by either a dual-SMAD method or a direct lineage reprogramming method. We found that a co-culture setting was necessary in maturing both the hiPSC-derived neurons and astrocytes. Besides, we showed that the neuron-astrocyte co-culture model was suitable for modeling the astrocyte-driven neuroinflammation. Overall, these hiPSC-derived neural cells represent powerful tools for both basic and translational studies.

## Introduction

*In-vitro* neural models are powerful tools in dissecting neurological phenomena at cellular and molecular level, and can provide a high-throughput drug screening platform. Neural cells from various animals have been widely used. However, failures in clinical translation of drug candidates indicated substantial species gaps in the neurophysiology between animals and humans [1–4]. Accessibility to human neural cells grants better authenticity in modeling human brain physiology and in pre-clinical evaluation of therapeutical molecules [5]. However, primary human neural cells are not easily accessible, and their donor-to-donor variance could share limited reproducibility. Differentiation of human induced pluripotent stem cells (hiPSCs) has emerged as a powerful way to derive the neural cells. Neural cells derived from hiPSCs can be produced in large scale and are found to closely resemble the primary ones [6]. This makes the hiPSC-neural cells an ideal alternative to the use of expensive primary cells or unfaithful immortalized cell lines. Here, two distinct methods were explored to generate neurons and astrocytes from hiPSCs. As a classic and well-developed method, we first used the dual-SMAD inhibition protocol [7] to generate midbrain floor plate precursor (mFPP)-derived midbrain dopaminergic (mDA) neurons and astrocytes. Next, we generated the same cell types using a novel approach known as direct lineage reprogramming, which is mediated by overexpression of lineage-restricted transcriptional factors (TFs). Both the mDA neurons and astrocytes from the two methods were found to express cell-type specific genes. We further showed that a co-culture setting was necessary for the maturation of both neural types. Moreover, we showed that our co-culture model is suitable for studying astrocyte-induced neuroinflammation. This model can be of valuable interest for basic and translational purposes.

### Methods Cell culture

HiPSCs were generated from IMR90 fibroblast cell line by episomal plasmid transfection method as previously reported [8]. The hiPSCs were maintained feeder-free in StemFlex medium (A3349401, Thermo Fisher) in 6-well plates coated with hESC-qualified Matrigel (#354277, Corning) in 5% CO_2_ humidified incubator at 37 ℃ and medium was refreshed daily. Upon reaching 70% confluency, cells were detached with ReleSR (#05872, STEMCELL) and sub-cultured as colonies at the ratio of 1:10- 1:20 in new coated well plates. HiPSCs were used for experiments no higher than passage 50.

Primary human astrocytes (#1800, ScienCell) were maintained in culture flasks coated with hESC-qualified Matrigel in 5% CO2 humidified incubator at 37 ℃. AGM medium (#1801, ScienCell) supplemented with 1 × Antibiotic Antimycotic (A5955, Sigma-Aldrich) was used for primary astrocyte culture. Medium was refreshed every two days. Upon reaching confluency around 90%, cells were detached with 0.025% trypsin EDTA (#25300054, Thermo Fisher) and sub-cultured at the density of 5,000 cells/cm^2^ in new coated flasks. Cells were used for experiments before passage 10.

HEK 293T cells (CRL11268, ATCC) were maintained in culture flasks in 5% CO_2_ humidified incubator at 37 ℃. HEK medium consists of Dulbecco’s Modified Eagle Medium (DMEM, #10313021, Thermo Fisher) supplemented with 10% (v/v) fetal bovine serum (FBS, #12103C, Sigma-Aldrich) and 1 × Antibiotic Antimycotic. Upon reaching confluency of 80-90%, cells were detached with 0.025% trypsin EDTA and sub-cultured at ratio of 1:10-1:20 in new flasks. All cell lines were regularly tested for mycoplasma contamination and were all found negative.

### Plasmids

All lentiviral plasmids were either originally purchased from Addgene or previously cloned in our lab (**Table S1**). All plasmids were amplified using Plasmid Maxiprep kit (Sigma-Aldrich, NA0310). Plasmid elutes were mixed with 1/10_th_ the volume of 3 M sodium acetate (pH 5.5) followed by mixing with 3 times the volumes of 95% (v/v) cold ethanol (stored at -20°C). Next, the solution was incubated at -20°C for 1 h followed by centrifuge at 5000 × g at 4°C for 15 min. Plasmid pellets were washed once with 75% cold ethanol (stored at -20°C) and centrifuged again. Finally, the plasmid pellets were dried at room temperature (RT) for 1 h followed by resuspending in small volume (∼300 μl) of nuclease-free water (AM9938, Thermo Fisher).

### Lentivirus production

A total of 6.5 × 10^6^ HEK 293T cells were seeded in the 14.5 cm dishes (Greiner Bio-One, 639160) and cultured overnight. Next day, the medium was switched to high glucose DMEM medium (22.5 ml) 2 h before transfection. For transfection in one 14.5 cm dish, DNA mix consisting of TF/rtTA plasmid (32 μg), pRSV-Rev (6.25 μg), pMD2.G (9 μg) and pMDLg (12.5 μg), was prepared in sterile Milli-Q water and levelled to 1.05 ml in a 15 mL falcon tube. The DNA mix was then mixed with 2 M CaCl_2_ (150 μl) and incubated at RT for 15 min. Next, 1.3 ml of 2 × HBS buffer (pH 7.4), which consisted of 281 mM NaCl, 100 mM HEPES, 1.5 mM Na_2_HPO_4_, was dropwise mixing with the DNA-calcium mix on a vortex (∼2000 rpm). After mixing, the transfect solution was immediately added dropwise to the HEK cells across the whole dish followed by gently swirling the dish to evenly distribute the transfectant. After 18 h, cells were fed with fresh high glucose DMEM (no less than 16 ml) and were kept in the incubator for 36-48 h. Medium containing viruses was collected and ultracentrifuged at 20,000 × g for 2 h at 20 °C. The virus pellets were resuspended in small amount PBS and stored at - 80 °C.

### Virus validation

Lentiviruses were titrated using an ELISA-based Lenti Titer Kit (TR30038, ORIGENE). The viruses were then validated in cultured cells. HEK 293T cells were seeded in 6-well plates at the density of 3 × 10^4^ cells/cm^2^. Next day, the cells were infected with individual TF viruses along with the rtTA virus, both of which were used at multiplicity of infection (MOI) of 10. Next day, the cells were washed twice with fresh medium to get rid of any residual viruses. The cells were then cultured in the presence of 1 μg/ml doxycycline (DOX, D3447, Sigma-Aldrich) in the medium for 48 h to induce expression of the TF. Total RNA of the cells was isolated and subjected to quantitative polymerase chain reaction (qPCR) to check expression of the TF.

### RNA isolation and qPCR

Cells growing at high density in the well plates were detached by trypsin or Accutase, washed and pelleted. Total RNA of the cells was isolated using ReliaPrep™ RNA isolation kit (Z6011, Promega) followed by cDNA synthesize from 1 μg RNA using IScript™ cDNA Synthesis Kit (BIO-RAD, 1708890). QPCR was performed using ITaq™ Universal SYBRR Green Supermix RT-qPCR Kit (BIO-RAD, 172-5124) and CFX96 Real-Time PCR System C1000™ Thermal Cycler (BIO-RAD). For data analysis, the cycle threshold (Ct) values of each gene were processed by 2^(-ΔΔCt)^ method to demonstrated fold-change in the gene expression [9]. All qPCR primer sequences can be found in **Table S2**.

### Differentiation of dopaminergic neurons from mFPPs

A schematic representation of the protocol can be found in **Figure 1a**. Differentiation of hiPSCs into mFPPs and subsequently into dopaminergic neurons was adapted from a published protocol with few modifications [10]. Briefly, hiPSC culture at ∼70% confluency was washed once with PBS and incubated with Accutase (A6964, Sigma-Aldrich) for 10 min in the incubator followed by pipetting for less than 10 times to generate homogeneous single cells. Cells were pelleted for 5 min at 300 × g, resuspended in StemFlex medium supplemented with 10 µM Y27632 (#72302, STEMCELL), and seeded in Matrigel coated 6-well plates at the density of 80,000 cells/cm^2^. Upon confluency, cells were patterned towards the mFPP identity by exposure of purmorphamine (#72202, STEMCELL), SB431542 (#616461, Sigma-Aldrich), CHIR99021 (SML1046, Sigma-Aldrich), SHH (#464-SH, R&D systems), LDN193189 (#72147, STEMCELL) and FGF-8 (#100-25, Peprotech) in the N2 medium which basically consisted of 1:1 mixed DMEM/F12 (#11320033, Thermo Fisher) and Neurobasal (#21103049, Thermo Fisher), supplemented with 1 × N2 supplement (#17502048, Thermo Fisher), 1 × GlutaMAX (#35050061, Thermo Fisher). Medium was refreshed daily. From day 11 onwards, medium was switched to the B27 medium which basically consisted of Neurobasal, supplemented with 1 × B27 supplement (#12587010, Thermo Fisher), 1 × GlutaMAX, 0.5 × non-essential amino acids (NEAA, #11140050, Thermo Fisher), 0.5 × sodium pyruvate (#11360039, Thermo Fisher) and 1 × Antibiotic Antimycotic. Medium was refreshed every two days. On day 11 and day 16, confluent cell layer was detached using Accutase and re-plated with 10 µM Y27632 at the density of 8 × 10^5^ cells/cm^2^ in Matrigel coated well plates. On day 25, cells were re-plated with 10 µM Y27632 at the density of 1 × 10^5^ cells/cm^2^ in PLO/LN-111 dual-coated well plates for final differentiation. For dual-coating, well plate bottom was first coated with 0.01% (v/v) Poly-L-ornithine (PLO, P4957, Sigma-Aldrich) solution at RT for at least 1 h followed by 3 times washing with Milli-Q water. The plates were then coated with 20 μg/ml Laminin-111 (LN111, BioLamina) in Ca^2+^/Mg^2+^ containing PBS (#14040141, Thermo Fisher) in the incubator for at least 2 h. Neurons were differentiated for up to 80 days. For culture additives, 2 µM purmorphamine was given to the cells from day 0 to day 6; 1.3 µM CHIR99021, 10µM SB431542, 100 nM LDN193189 and 200 ng/ml SHH from day 0 to day 9; 100 ng /ml FGF-8 from day 9 to day 16; 20 ng/ml BDNF (#450-02, Peprotech), 20 ng/ml GDNF (#450-10, Peprotech) and 0.2 mM ascorbic acid (A4403, Sigma-Aldrich) from day 11 to day 80; 0.5 mM db-cAMP (D0627, Sigma-Aldrich), 10 µM DAPT (#2634, TOCRIS) and 1 ng/ml TGF-β3 (#100-36E, Peprotech) from day 22 to day 80. Immunofluorescence microscopy (IFM) was performed on day 45 whereas qPCR was performed on day 80.

**Figure 1.**
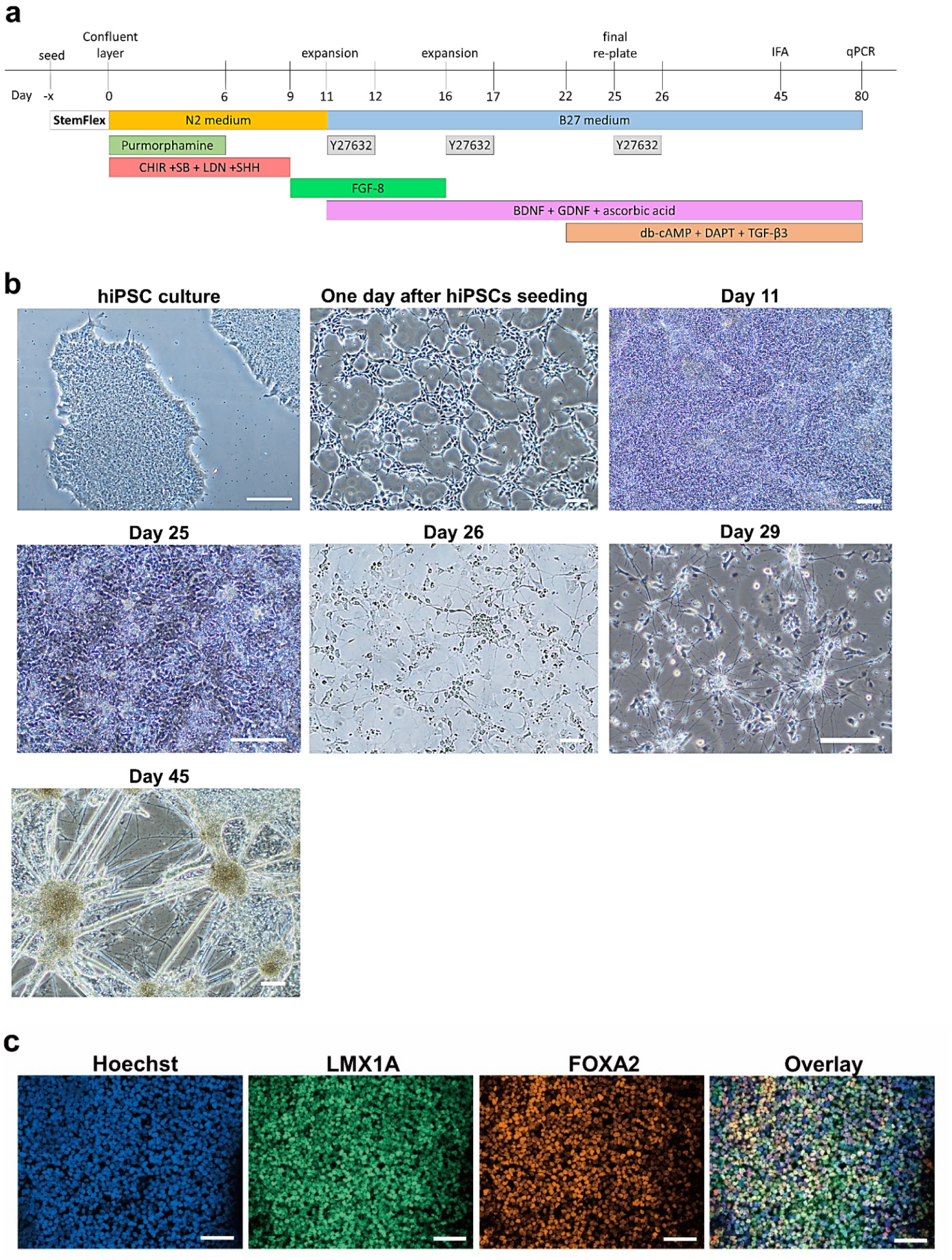
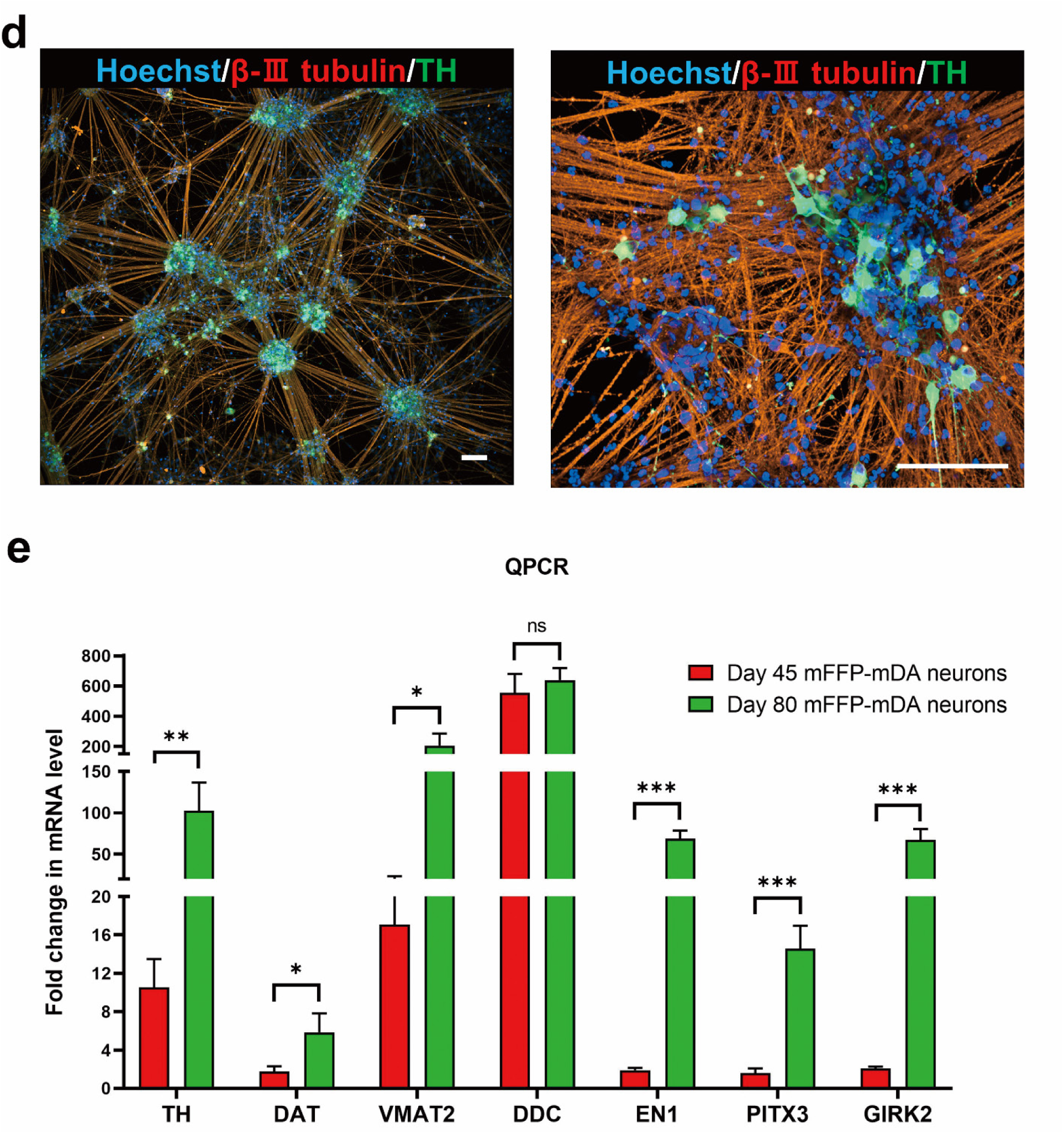
Differentiation of hiPSCs into mFPPs and subsequently into mDA neurons. **(a)** Schematic illustration of the differentiation protocol. **(b)** Brightfield images during mDA neuron differentiation. Scale bars are 200 µm. **(c)** IFM images showing the day 11 mFPPs. Scale bars are 50 µm. **(d)** IFM images showing the day 45 mDA neurons. Scale bars are 100 µm. **(e)** QPCR quantification of the mDA neurons. HiPSCs were used as control cells (not shown). Graph shows mean ± SD. Statistical analysis was performed using student’s t-test; ns indicates no significant differences; \**P* < 0.05, \*\**P* < 0.01, \*\*\**P* < 0.001.

### Differentiation of astrocytes from mFPPs

A schematic representation of the protocol can be found in **Figure 2a**. The hiPSC culture was patterned towards mFPPs as mentioned above. For the differentiation of astrocytes from mFPPs, the protocol was adapted from two published papers with few modifications [11, 12]. Briefly, from day 11 to day 21, the NPC medium was used which basically consisted of Neurobasal, supplemented with 0.5 × N2 supplement, 0.5 × B27 supplement and 1 × GlutaMAX. Medium was refreshed every two days. On day 11, confluent cell layer was detached using Accutase and re-plated with 10 µM Y27632 at 1:2 ratio in Matrigel coated well plates. On day 15, confluent cell layer was detached again and re-plated at 1:4 ratio. On day 21, cells were re-plated for final differentiation at the density of 1.5 × 10^4^ cells/cm^2^ in Matrigel coated well plates. Next day, the medium was switched to AM medium. Astrocytes were differentiated for up to 52 days. Medium was refreshed every two days. Once the confluency reached ∼90%, cells were to be re-plated at the density of 1.5 × 10^4^ cells/cm^2^ in Matrigel coated well plates. For culture additives, 2 µM purmorphamine was given to the cells from day 0 to day 6; 1.3 µM CHIR99021, 10µM SB431542, 100 nM LDN193189 and 200 ng/ml SHH from day 0 to day 9; 20 ng /ml FGF-2 (#100-18B, Peprotech) and 20 ng/ml EGF (AF-100-15, Peprotech) from day 11 to day 21. IFM was performed on day 45 whereas qPCR was performed on day 52.

**Figure 2.**
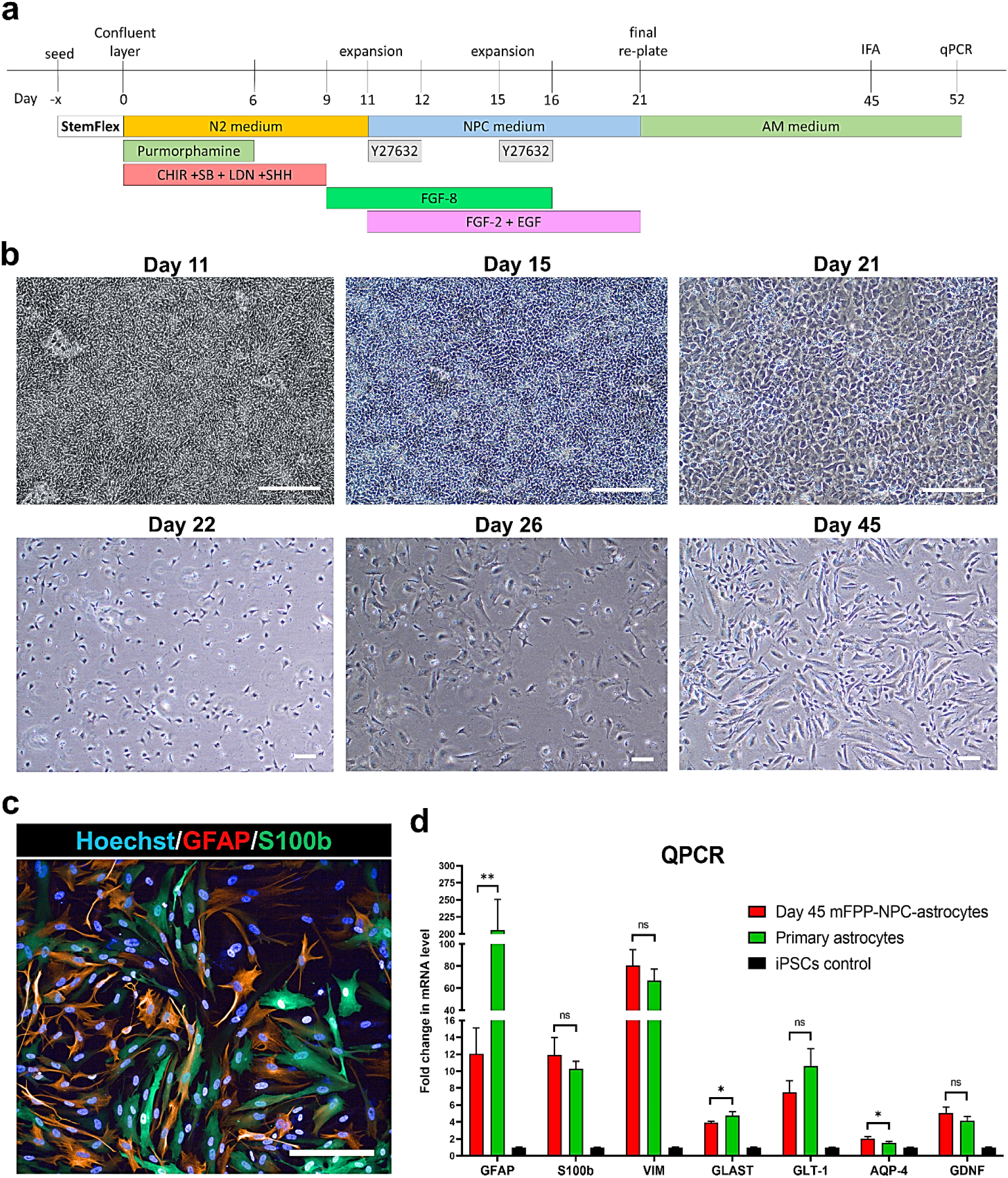
Differentiation of hiPSCs into mFPPs and subsequently into astrocytes. **(a)** Schematic illustration of the differentiation protocol. **(b)** Brightfield images during astrocyte differentiation. Scale bars are 200 µm. **(c)** IFM images showing the day 45 astrocytes. Scale bar is 200 µm. **(d)** QPCR quantification of the day 52 astrocytes. Graph shows mean ± SD. Statistical analysis was performed between generated astrocytes and primary astrocytes using student’s t-test; ns indicates no significant differences; \**P* < 0.05, \*\**P* < 0.01.

### Differentiation of dopaminergic neurons from hiPSCs by direct lineage reprogramming

A schematic representation of the protocol can be found in **Figure 3a**. Differentiation of hiPSCs into induced dopaminergic (DA) neurons was adapted from a published protocol with few modifications [13]. Briefly, single hiPSCs were seeded in StemFlex medium supplemented with10 µM Y27632 in Matrigel coated 6-well plates at the density of 80,000 cells/cm^2^. Next day, cells were infected with 4 lentiviruses (rtTA, ASCL1, LMX1A, NURR1, each at MOI=10) in fresh StemFlex medium supplemented with 10 µM Y27632 and 8 μg/ml polybrene (Merck, TR-1003-G). The infection lasted for 17 hours. After infection, fresh StemFlex medium supplemented with 1 µg/ml DOX was used to induce expression of the TFs. This day was marked as day 0 of differentiation. From this moment, DOX was consistently presented in the medium until day 14. From day 1 to day 7, cells were daily fed with the N2 medium which basically consisted of DMEM/F12, supplemented with 1 × N2 supplement, 1 × GlutaMAX and 0.5 × NEAA. Cells were selected by 0.125 µg/ml puromycin from day 1 to day 5 (**Figure S1a left**). On day 6, cells were detached with Accutase, pelleted, resuspended in 10 µM Y27632 supplied N2 medium and re-plated in PLO/LN-111 dual-coated well plates at the density of 1 × 10^5^ cells/cm^2^. The day after re-plate, medium was switched to the B27 medium which basically consisted of 1:1 mixed DMEM/F12 and Neurobasal, supplemented with 1 × B27 supplement, 1 × GlutaMAX, 0.5 × NEAA, 0.5 × sodium pyruvate and 1 × Antibiotic Antimycotic. Medium was changed every two days. DA neurons were differentiated for up to 28 days. For culture additives, 0.2 µM ascorbic acid was given to the cells from day 1 to day 28; 0.5 mM db-cAMP, 20 ng/ml BDNF, 20 ng/ml GDNF from day 5 to day 28; 10 µM DAPT from day 7 to day 28. IFM was performed on day 21 whereas other assays were performed on day 28.

**Figure 3.**
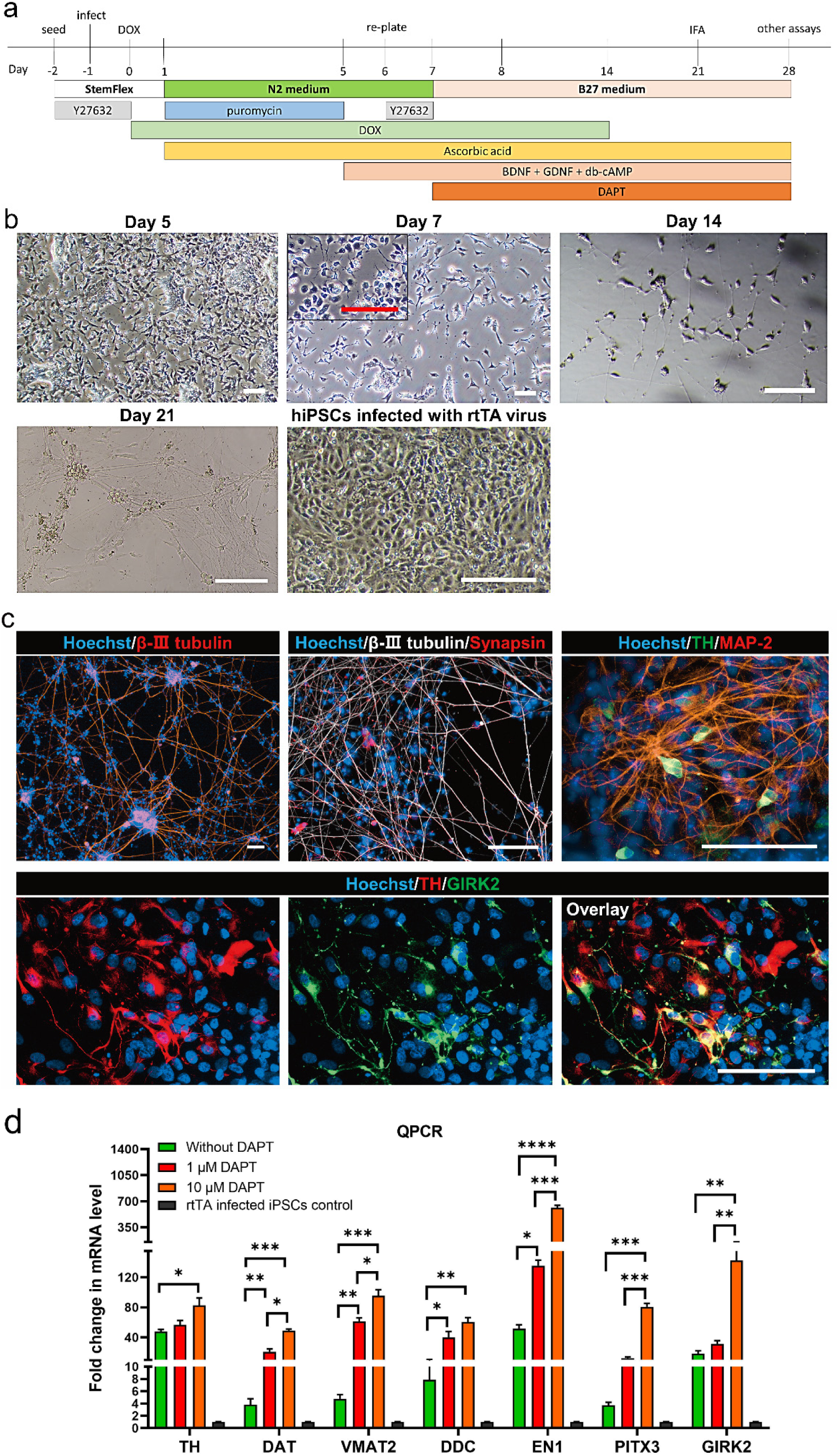
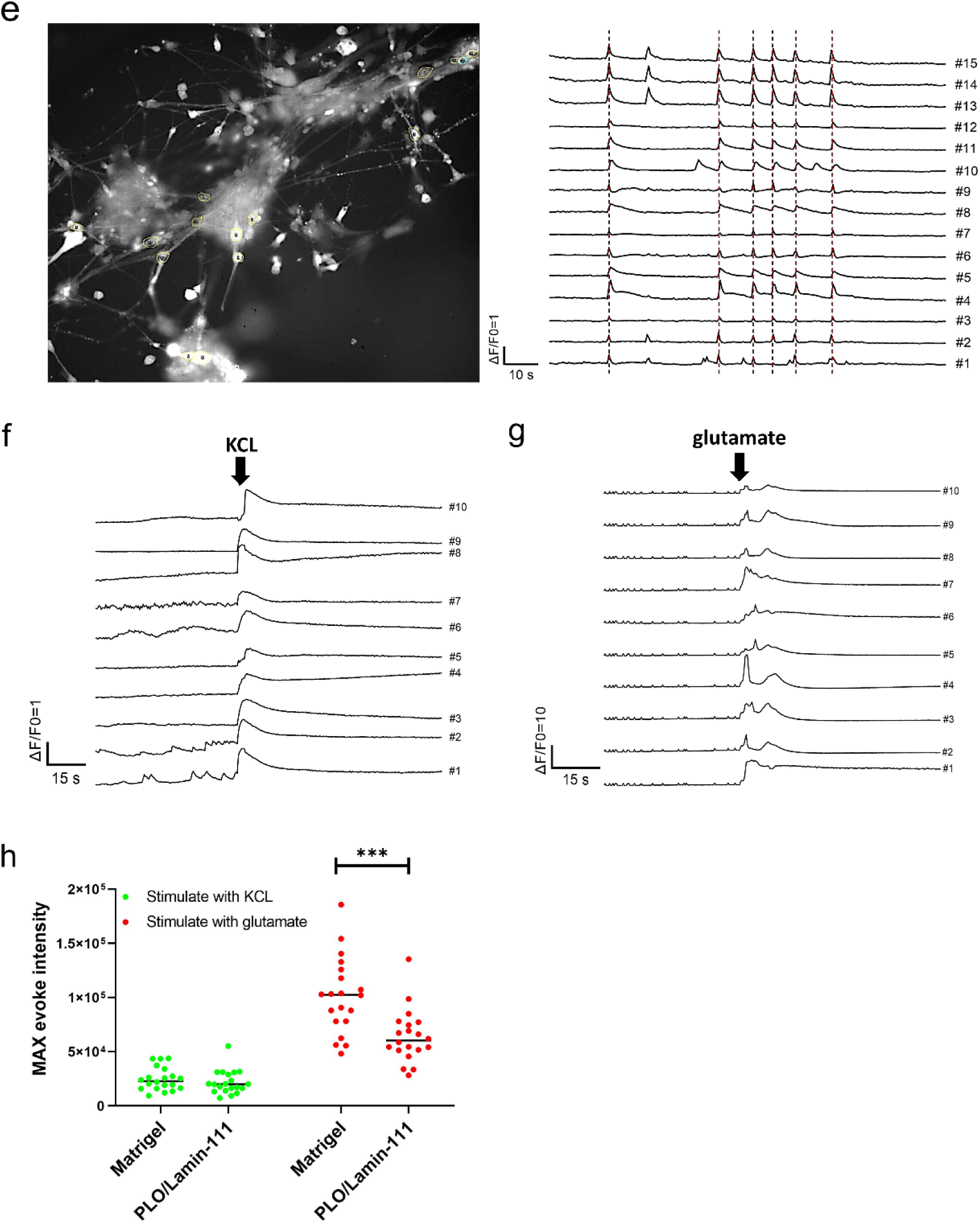
Differentiation of hiPSCs into ALN-neurons by direct lineage reprogramming. **(a)** Schematic illustration of the differentiation protocol. **(b)** Brightfield images during the ALN-neuron differentiation. Scale bars are 200 µm. **(c)** IFM images showing the day 21 ALN-neurons. Scale bars are 100 µm. **(d)** QPCR quantification of the day 28 ALN-neurons. The rtTA-infected hiPSCs were used as control cells. **(e)** Calcium imaging showing basal signals in the day 28 ALN-neurons. Signals of 15 selected neurons with synchronized signaling were plotted. **(f, g)** Calcium imaging of the day 28 ALN-neurons in response to KCL **(f)** and glutamate **(g)** stimulations. For each group, the signals of 10 selected neurons were recorded. **(h)** Calcium imaging of the day 28 ALN-neurons cultured on different coating materials. Each dot represented a cell. All graphs show mean ± SD. Statistical analysis was performed using one-way ANOVA followed by Dunnett’s multiple comparisons tests for **(d)** and using student’s t-test for **(h)**; ns indicates no significant differences; \**P* < 0.05, \*\**P* < 0.01, \*\*\**P* < 0.001.

### Differentiation of astrocytes from hiPSCs by direct lineage reprogramming

A schematic representation of the protocol can be found in **Figure 4a**. Differentiation of hiPSCs into induced astrocytes was adapted from a published protocol with few modifications [14]. Briefly, single hiPSCs were seeded in StemFlex medium containing 10 µM Y27632 in Matrigel coated 6-well plates at the density of 60,000 cells/cm^2^. Next day, cells were infected with 4 lentiviruses (rtTA, NFIA, NFIB, SOX9, each at MOI=10) in fresh StemFlex medium supplemented with 10 µM Y27632 and 8 μg/ml polybrene. The infection lasted for 17 hours. After infection, fresh StemFlex supplemented with 1 µg/ml DOX was used to induce expression of the TFs. This day was marked as day 0 of differentiation. From this moment, DOX was consistently presented in the medium until day 14. The next two days, cells were daily fed with the expansion medium which basically consisted of DMEM/F12, supplemented with 1 × N2 supplement, 1 × GlutaMAX and 5% FBS. Cells were selected by 0.2 µg/ml puromycin from day 1 to day 5 (**Figure S1a right**). From day 3-6, medium was gradually switched to the FGF medium which basically consisted of Neurobasal, supplemented with 2% FBS, 1 × B27 supplement and 1 × NEAA. On day 7, cells were detached using Accutase, pelleted, resuspended and re-plated in new Matrigel coated well plates at the density of 5 × 10^4^ cells/cm^2^. From day 8, the astrocyte maturation medium was used which basically consisted of 1:1 mixed F12/DMEM and Neurobasal, supplemented with 2% FBS, 1 × N2 supplement, 1 × GlutaMAX, 1 mM sodium pyruvate and 1× Antibiotic Antimycotic. Medium was changed every two days. Cells were differentiated for up to 28 days. Cells were re-plated again on day 21 in case of high confluency. For culture additives, 0.2 mM ascorbic acid was given to the cells from day 1 to day 28; 5 ng/ml FGF-2 from day 3 to day 7; 5 ng/ml CNTF (#450-13, Peprotech) and 5 ng/ml BMP-4 (#120-05ET, Peprotech) from day 3 to day 21; 0.5 mM db-cAMP from day 7 to day 21; 5 ng/ml HB-EGF (#259-HE, R&D systems) from day 7 to day 28. IFM was performed on day 21 whereas other assays were performed on day 28.

**Figure 4.**
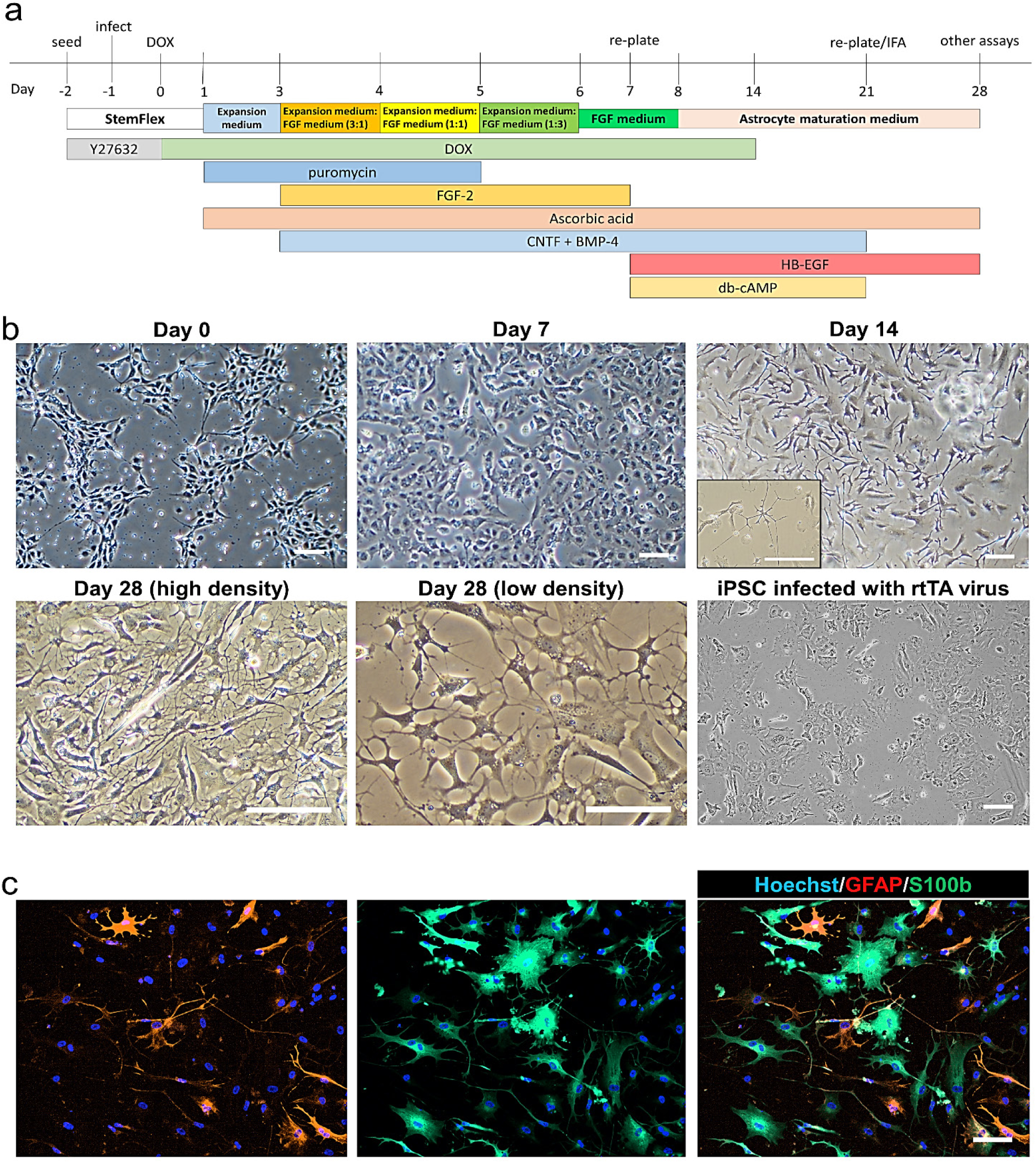
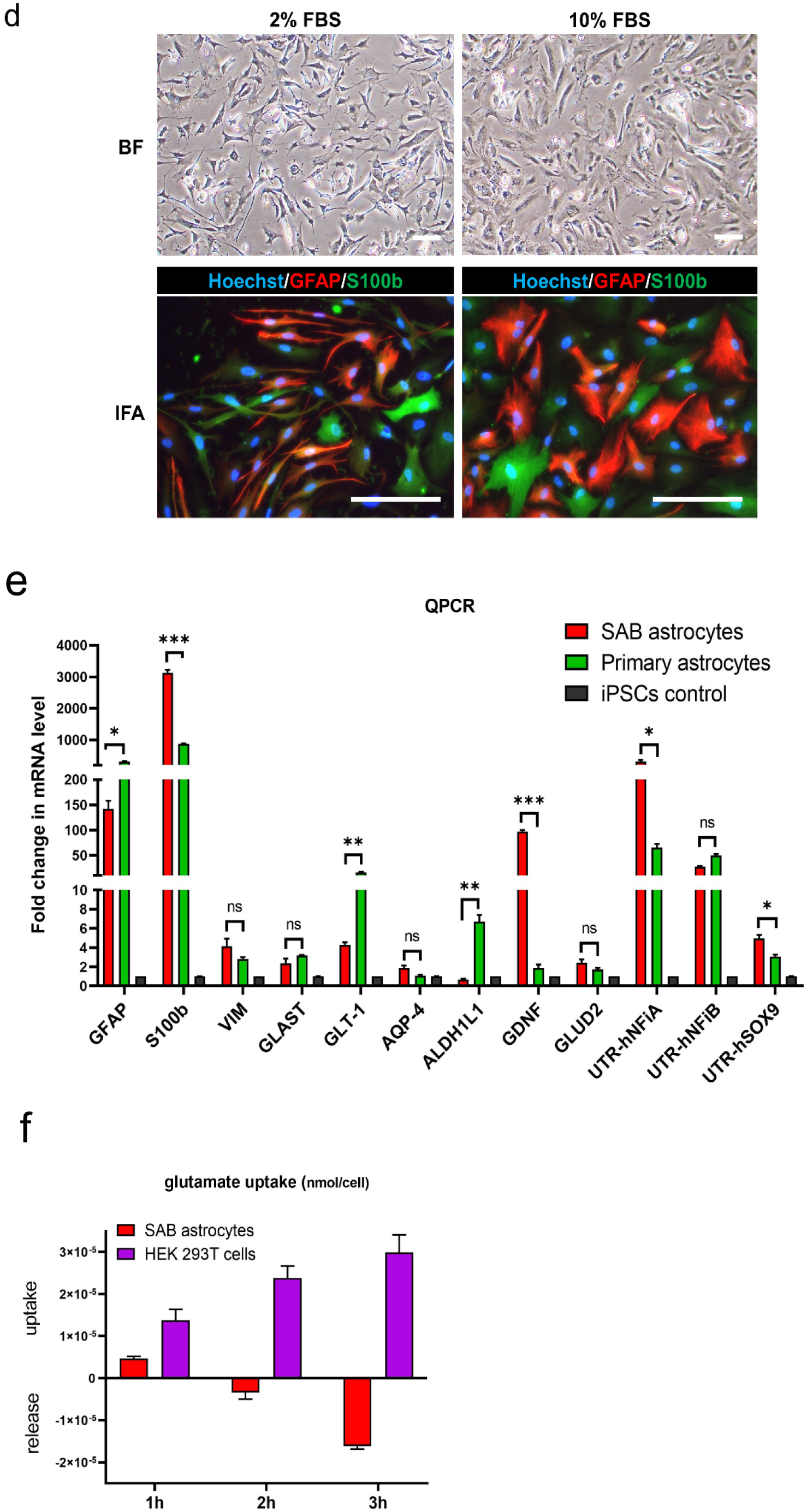
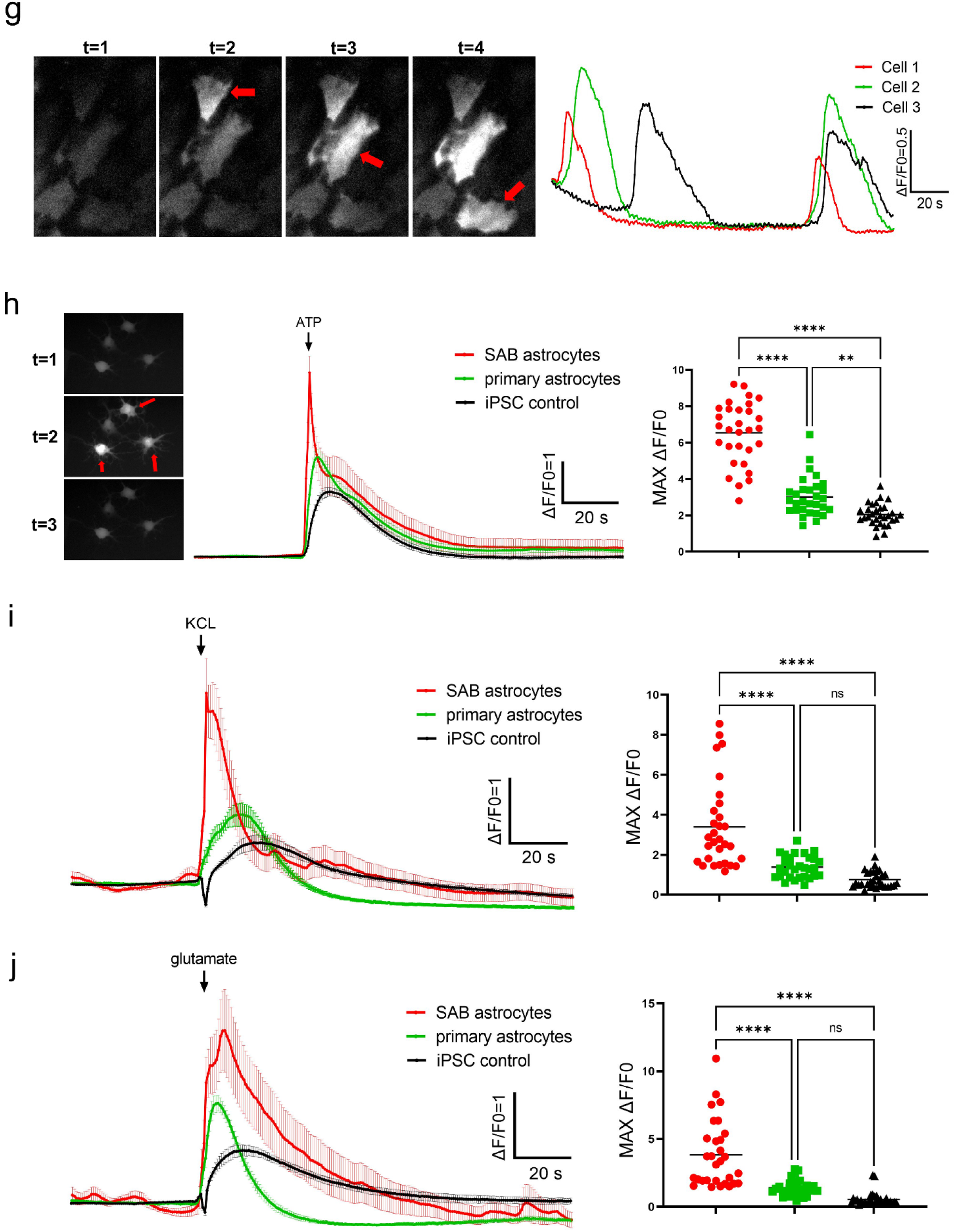
Differentiation of hiPSCs into SAB-astrocytes by direct lineage reprogramming. **(a)** Schematic illustration of the differentiation protocol. **(b)** Brightfield images during the SAB-astrocyte differentiation. Scale bars are 200 µm. **(c)** IFM images showing the day 21 SAB-astrocytes. Scale bars are 100 µm. **(d)** Brightfield and IFM images of the SAB-astrocytes pre-cultured in medium containing different FBS levels for 5 days. Scale bars are 200 µm. **(e)** QPCR quantification of the day 28 SAB-astrocytes. The rtTA-infected hiPSCs were used as control cells. **(f)** Glutamate uptake of the day 28 SAB-astrocytes. HEK 293T cells were used as control cells. **(g)** Calcium imaging showing basal signals in the day 28 SAB-astrocytes. **(h-j)** Calcium imaging of the day 28 SAB-astrocytes in response to ATP **(h)**, KCL **(i)** and glutamate **(j)** stimulations. For each group, 30 selected cells were recorded. All graphs show mean ± SD. Statistical analysis was performed between SAB-astrocytes and primary astrocytes using student’s t-test for **(e)** and using one-way ANOVA followed by Tukey’s multiple comparisons tests for **(h-j)**; ns indicates no significant differences; \**P* < 0.05, \*\**P* < 0.01, \*\*\**P* < 0.001.

### Astrocyte-neuron co-culture

A schematic representation of the co-culture protocol can be found in **Figure 5a**. For non-contact co-culture in the transwell system, membrane of the inserts (0.4 µm pore size, #140640, Thermo Fisher) was coated with Matrigel whereas bottom of the outer wells was coated with PLO/LN-111. The day 5 lentiviral-derived DA neurons were harvested and re-plated in the well bottom at the density of 1 × 10^5^ cells/cm^2^. Meanwhile, the day 7 lentiviral-derived astrocytes were harvested and re-plated in the inserts at the density of 1.6 × 10^5^ cells/cm^2^. This day was marked as day 0 of co-culture. For direct-contact co-culture in the well plates, neurons and astrocytes were co-seeded in the PLO/LN-111 dual-coated well plates at the density of 1 × 10^5^ cells/cm^2^ and 5 × 10^4^ cells/cm^2^, respectively. From day 0 to day 7, the co-culture N2 medium was used which basically consisted of 1:1 mixed F12/DMEM and Neurobasal, supplemented with 1 × N2 supplement, 1 × GlutaMAX, 0.5 × NEAA, 0.5 × sodium pyruvate and 1 × Antibiotic Antimycotic. From day 7 onwards, N2 supplement was replaced by B27 supplement in the medium, which subsequently termed as the co-culture B27 medium. Medium was refreshed every two days. For culture additives, 1 µg/ml DOX was given to the cells from day 0 to day 7; 1 µM DAPT from day 7 to day 21; 0.2 mM ascorbic acid, 0.5 mM db-cAMP, 10 ng/ml BDNF, 10 ng/ml GDNF from day 0 to day 21.

**Figure 5.**
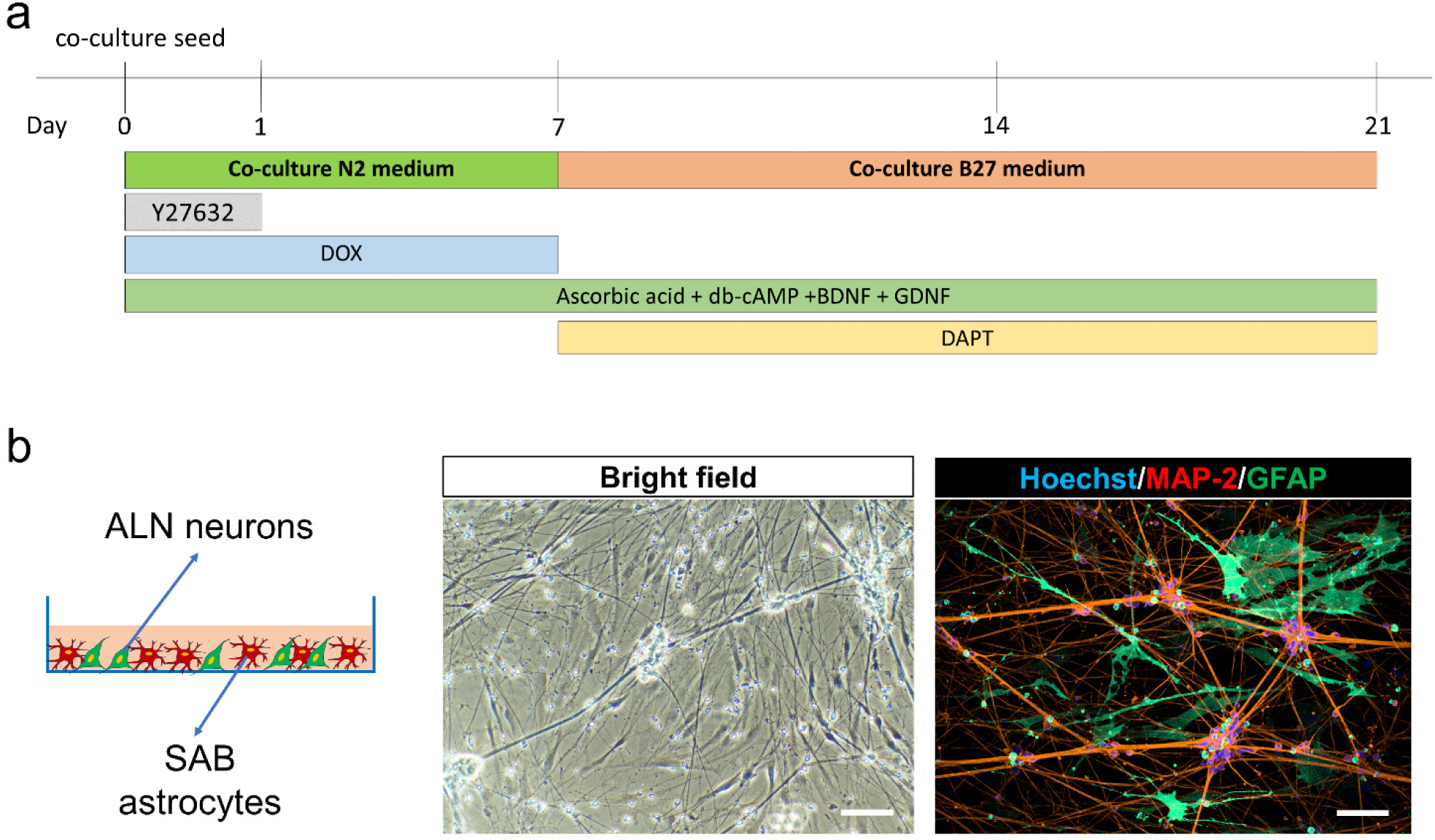
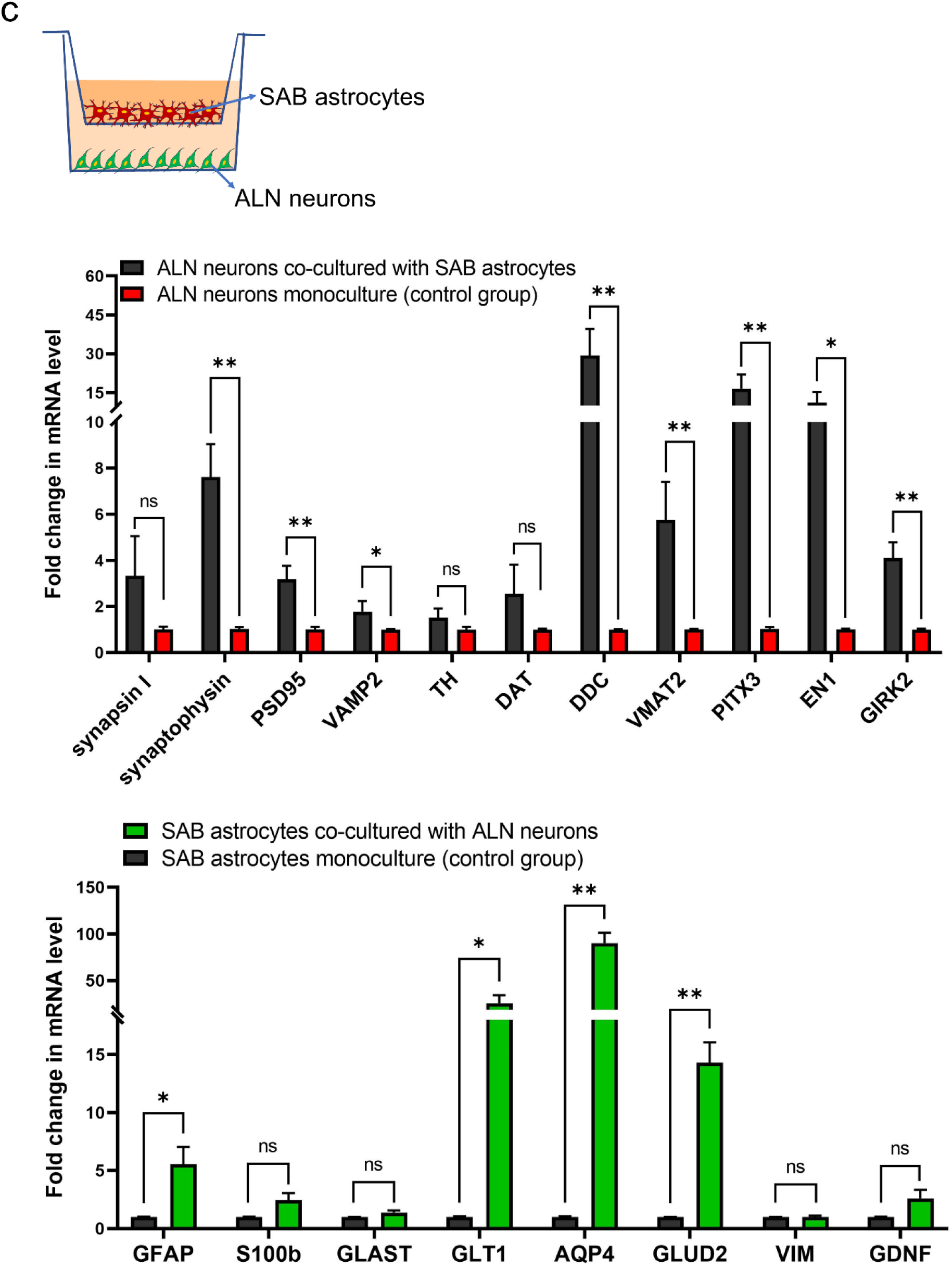
Co-culture model consisting of SAB-astrocytes ALN-neurons. **(a)** Schematic illustration of the co-culture seeding protocol. **(b)** Brightfield and IFM images showing the co-culture in a contactable manner (in well plates). Scale bars are 100 µm. **(c)** QPCR quantification of SAB-astrocytes and ALN-neurons when co-cultured in a non-contactable way (in transwell system). All graphs show mean ± SD. Statistical analysis was performed using student’s t-test; ns indicates no significant differences; \**P* < 0.05, \*\**P* < 0.01.

### Inflammatory astrocyte induction

This assay was performed in the transwell-based co-culture model. The day 5 lentiviral-derived DA neurons were seeded and differentiated in the outer well bottom of the transwell, as described earlier. To induce neuroinflammatory astrocytes generally known as A1 type [15], the day 28 lentiviral-derived astrocytes were incubated with 10 ng/ml TNFα (#300-01A, Peprotech) and 10 ng/ml IL-1α (#200-01A, Peprotech) in basal astrocyte maturation medium for 48 h in the 6-well plates. The astrocytes were then collected and re-plated in the inserts. The co-culture model was maintained for 3 days in additive-free co-culture B27 medium without medium refresh. Total RNA of the neurons and astrocytes were separately collected and analyzed by qPCR.

### Glutamate uptake

The lentiviral-astrocytes, primary astrocytes or HEK293T were seeded in Matrigel coated 6-well plates and cultured till confluency. A total of 1 ml glutamate (G1251, Sigma-Aldrich) working solution, which was prepared in HBSS buffer at the concentration of 100 μM, was given to the cells and incubated in the incubator for 3 h. Medium was then harvested, centrifuged to collect the supernatant. In parallel, cells remaining adhesive in the wells were collected for cell counting. Measurement of glutamate uptake was performed following instruction of the glutamate assay kit (ab138883, abcam).

### Calcium imaging

Lentiviral-derived astrocytes and neurons were re-plated in Matrigel or PLO/LN-111 coated 96-well plates. Assay medium was prepared by diluting Fluo-4 AM (ab241082, Abcam) dye in Neurobasal medium to make the final concentration of 5 µM. A total of 100 μl assay medium was given to the cells per well and incubated for 30 min in the incubator. Assay medium was then replaced with fresh Neurobasal medium. To detect spontaneous calcium fluctuations, cells were imaged under confocal microscope (YOKOGAWA, 20 × magnification, Ex/Em 494/506 nm) at 2 Hz for 2-3 min. To detect evoked cell response to stimuli, cells were stimulated with 50 μM ATP (FLAAS-1VL, Sigma-Aldrich)/65 mM KCL/100 μM glutamate using dispensing function of the microscope. For data processing, fluorescence intensity of the region of interest (ROI) was measured using ImageJ software and the profile was plotted [16].

### Immunofluorescence microscopy (IFM)

All IFM steps were carried out at RT unless specified otherwise. Cells growing in the well plates were washed twice in PBS and fixed in 4% (v/v) paraformaldehyde (PFA) for 10 min. Cells were then washed three times in PBS and permeabilized in PBS containing 0.1% (v/v) TritonX-100 (T8787, Sigma-Aldrich) for 10 min. After permeabilization, cells were blocked in PBS containing 2% (w/v) bovine serum albumin (BSA) for a minimum 1 h. Cells were incubated overnight at 4 °C with primary antibodies for S100β (S100 calcium-binding protein B, GA504, Dako, 1:400), GFAP (glial fibrillary acidic protein, #76214, Dako, 1:200), β-Ⅲ tubulin (#32-2600, Thermo Fisher, 1:800), MAP-2 (microtubule associated protein 2, m2323, Sigma-Aldrich, 1:500), TH (Tyrosine hydroxylase, #6211, Abcam, 1:1000), LMX1A (LIM homeobox transcription factor 1 alpha, #139726, Abcam, 1:500), FOXA2 (Forkhead box protein A2, sc-374376, SANTA CRUZ, 1:50), GIRK2 (G-protein-activated inward rectifying potassium type 2, #66502, Abcam, 1:200), Synapsin-1 (ab8, Abcam, 1:500) in blocking buffer. Next day, cells were washed three times with PBS, incubated with fluorescence labeled secondary antibodies including Goat anti-Rabbit Alexa Fluor-488 (A-11008, Thermo Fisher, 1:500), Goat anti-Rabbit Alexa Fluor-555 (A-21428, Thermo Fisher, 1:500), Goat anti-Mouse Alexa Fluor-568 (A-21422, Thermo Fisher, 1:500) in blocking buffer for 2 h in dark followed by another 10 min incubation with Hoechst (#62249, Thermo Fisher, 1:1000) in PBS to stain the nuclei. Cells were finally washed three times in PBS and images were acquired by confocal microscopy.

### Statistical analysis

All data were presented as mean ± standard deviation (SD). Statistical analysis was performed by student’s t-test or one way ANOVA method using GraphPad Prism 9 software. For multiple comparisons, either Tukey’s multiple comparisons tests or Dunnett’s multiple comparisons tests was performed. P values < 0.05 were considered statistically significant.

## Results

### Generation of dopaminergic neurons and astrocytes from mFPPs

Neural differentiation through the dual-SMAD inhibition approach has been well studied [7, 17]. We first explored this method to derive human neurons and astrocytes from hiPSCs. Confluent hiPSC monolayer was induced to neural fate by exposure to SB431542 and LDN193189. Specifically, we patterned the cells further to midbrain floor plate (mFPP) fate by co-activation of the Sonic hedgehog and Wnt/β-catenin pathways using purmorphamine, CHIR99021 and SHH. On day 11, differentiated cells appeared highly compact and homogeneous (**Figure 1b**). These mFPPs co-expressed LMX1A and FOXA2, two specific markers for floor plate identity (**Figure 1c**) [10]. We then expanded the mFPPs twice to deplete their proliferative potential and to further increase the mDA differentiation efficiency [18]. Final differentiation of the mDA neurons was induced by re-plating the expanded mFPPs from day 25 into PLO/Lamin-111 dual-coated wells at low density. To assist neuron maturation, several neurotrophic factors were supplied in the medium including BDNF, GDNF, ascorbic acid, db-cAMP, DAPT and TGF-β3. One day after re-plate, the neurons already showed neurite outgrowth (**Figure 1b**). By day 45, these neurons have formed extensive neural networks. Moreover, neuron clusters were formed, and these structures were interconnected by thick nerve bundles (**Figure 1b, d**). At the stage, the β-III tubulin positive neurons were found to co-express TH, a rate-limiting enzyme in the biosynthesis of dopamine (**Figure 1d**). This indicated the commitment of the dopaminergic fate of the neurons. For further maturation, the neurons were cultured for up to 80 days. Compared to day 45, these neurons were further enriched for a number of mDA markers including DAT, VMAT2, DDC, EN1, PITX3 (**Figure 1e**). Importantly, GIRK2, a nucleus A9 specific marker in the ventral midbrain, was expressed by the neurons. This indicated that the protocol was able to generate the mDA neuron subtype which was specifically lost during Parkinson’s disease (PD) [19].

For the generation of astrocytes, the day 11 mFPPs were maintained and expanded in the NPC medium. Final differentiation of the astrocytes was induced by re-plating the expanded neural progenitor cells from day 21 into Matrigel coated wells at low density. Cells were matured in a commercial astrocyte medium with regular cell passage. On day 45, these cells adopted a fibroblast-like morphology (**Figure 2b**). IFM showed that the differentiated cells expressed astrocytic markers GFAP and S100β (**Figure 2c**). On day 52, more astrocytic genes were found positive in these differentiated astrocytes such as VIM, GLAST, GLT-1, AQP-4 and GDNF (**Figure 2d**). The expression of several genes became rather comparable to the primary astrocytes, indicating maturation of these hiPSC-derived astrocytes.

### Generation of dopaminergic neurons and astrocytes by direct lineage reprogramming

Recently, direct lineage reprogramming has become a novel approach to generate various cell types from the iPSCs [20]. In this section, we explored the possibility to derive mDA neurons and astrocytes by direct reprogramming. Lentiviruses were used to infect the hiPSCs to drive constant expression of the exogenous genes. To tightly control the expression of exogenous TFs, the lentiviruses were constructed under a drug-inducible expression system called Tet-On (**Figure S1b**) [21]. For this system, an additional virus expressing the rtTA is used, which functions as a reverse tet-transactivator. In the presence of DOX, rtTA binds to the Tet-On element and drives the expression of downstream genes.

For the generation of mDA neurons, single hiPSCs were infected with 4 viruses carrying ASCL-1, NURR1, LMX1A and rtTA, the former 3 were collectively referred to as ALN. DOX was given to the cells the day after infection to initiate TFs expression. Since the ASCL-1 virus carries a puromycin resistance gene, we were therefore able to select the cells successfully infected at least by ASCL-1, a critical regulator of neurogenesis. Several MOIs were tested to screen for the highest differentiation efficiency. As MOI increased, more infected hiPSCs were found to survive the selection (**Figure S1c**). Based on the results, MOI=10 was chosen for the experiment. After selection, the cells were collected and re-plated into PLO/Lamin-111 dual-coated wells. A clear neural morphology began to appear one day after the re-plate, where the cells developed branched neurite-like structures (**Figure 3b**). By day 14, the differentiated neurons could already form axonal connections. On day 21, more complex neural networks were seen, where the neurons spontaneously formed multicellular clusters that were interconnected by thick nerve tracks (**Figure 3b, c**). In comparison, the rtTA-infected hiPSCs did not show signs of neurogenesis. The ALN-neurons were positive for β-Ⅲ tubulin and also expressed TH, synapsin and GIRK2 (**Figure 3c**). The ALN-neurons showed expression of more mDA specific genes by day 28 including DAT, VMAT2, DCC, EN1, PITX3, indicating the adaptation of a midbrain dopaminergic subtype (**Figure 3d**). In addition, double-positive staining for TH and GIRK2 in some neurons suggested the presence of nucleus A9 subtype DA neurons in the culture. Inhibition of the Notch signaling pathway is shown to promote and accelerate neural differentiation and maturation [22, 23]. We found that the use of DAPT, a Notch inhibitor, significantly increased the expression of the abovementioned genes, indicating that the ALN-neurons reached a more matured stage (**Figure 3d**). Intracellular calcium waves are associated with a number of neural activities [24]. We confirmed the presence of calcium waves in the ALN-neurons, which indicated spontaneous neural activities (**Figure 3e**). Moreover, calcium signals could be synchronized among certain neurons. This meant the neural signals have propagated through neural synapses in a functional neural network. Besides, the ALN-neurons were able to respond to stimuli such as KCL and glutamate by eliciting an intracellular calcium wave (**Figure 3f, g**). Some early study has found that the coating material could affect neural activities [25]. Here, we found that the glutamate stimulation induced a significantly higher calcium intensity in neurons that grew on the PLO/Lamin-111 coating as compared to that grew on Matrigel coating (**Figure 3h**). However, neurons’ response to KCL was not affected in such ways.

For the generation of astrocytes, similarly, single hiPSCs were infected with 4 viruses carrying SOX9, NFIA, NFIB and rtTA, the former 3 were collectively referred to as SAB. Puromycin selection was done on the infected hiPSCs where SOX9 virus granted puromycin resistance. A clear astrocyte morphology began to appear on day 14 of differentiation (**Figure 4b**). By day 28, more complex multi-branches or star-shape morphology could be seen (**Figure 4b**). In comparison, the rtTA-infected hiPSCs only showed a flattened cell shape. The SAB-astrocytes were found to express two astrocytic markers GFAP and S100β (**Figure 4c**). It is known that astrocyte morphology can be influenced by the serum level. We cultured SAB-astrocytes in either a low FBS (2%) medium or a high FBS (10%) medium and found that astrocytes growing in high serum condition developed a more flattened morphology in 5 days, indicating a reactive status [2] (**Figure 4d**). The SAB-astrocytes showed expression of more astrocytic genes by day 28 including VIM, GLAST, GLT-1, AQP-4, ALDH1L1, GDNF, GLUD2 (**Figure 4e**). Comparison with primary astrocytes revealed lower levels for some of the genes in SAB-astrocytes such as GFAP, S100β, GLT-1 and ALDH1L, indicating possibly a less matured status of these astrocytes at the moment. In addition to ectopic overexpression, we also measured the expression of endogenous NFIA/NFIB/SOX9. By analyzing the untranslated region (UTR), it is possible to differentiate gene expressions between internal and ectopic sources. We found that endogenous NFIA/NFIB/SOX9 could be abundantly detected since DOX withdrawal from day 14 of differentiation (**Figure 4e**). This indicated that the cell fate towards astrocyte differentiation became irreversible for the SAB-infected hiPSCs at this moment.

One of the functions of astrocytes is to timely recycle excessive glutamate in the synaptic cleft to avoid neuron excitotoxicity [26]. We therefore tested the ability of the SAB-astrocytes to uptake glutamate. HEK293T cells were used as control since they were found to uptake glutamate to a subtle extent [27]. Despite that glutamate uptake was detected for SAB-astrocytes in the first hour, an unexpected glutamate release dominated for the following two hours (**Figure 4f**). Interestingly, primary astrocytes also showed a glutamate release profile in the assay (**Figure S2**). Notably, as astrocytes were found also to release glutamate and glutamine [26, 28], we speculated that the glutamate release, somehow triggered in the astrocytes, might overcome the uptake. Similar to neurons, SAB-astrocytes were found to exhibit spontaneous calcium oscillations and these calcium waves could spread among connecting astrocytes (**Figure 4g**), which was in line with other studies [11, 27, 29]. Notably, the speed of the calcium wave propagating among astrocytes was found to be delayed and slower compared to that crossing the neuron network. The SAB-astrocytes also responded to stimuli such as ATP, KCL and glutamate. We showed that the response patterns to these stimuli could readily differentiate among the SAB-astrocytes, primary astrocytes and hiPSC control. All three stimuli could evoke a sharp calcium oscillation in both the SAB-astrocytes and primary astrocytes, whereas the rtTA-infected hiPSCs only showed a gentle and weak reaction (**Figure 4h, i, j**). The amplitude of calcium spikes was significantly higher in the SAB-astrocytes than in the primary astrocytes, indicating a more active status of the SAB-astrocytes.

Taken together, we showed that both functional astrocytes and neurons can be generated from hiPSCs by overexpressing lineage-specific TFs.

### Astrocyte and neuron co-culture

Astrocytes assist in maturating the lineage reprogrammed neurons under *in-vitro* condition [14, 30–32]. We established a co-culture model, which comprised the SAB-astrocytes and ALN-neurons, in either a direct-contact format (**Figure 5b**) or in a non-contact format (**Figure 5c**). The direct-contact model allows for convenient spectating of neural morphology or key gene expressions under fluorescence microscope whereas the transwell-based non-contact model allows for studying how the co-culture condition influences individually each cell type.

We found that the co-culture environment mutually promoted the maturation of both the ALN-neurons and SAB-astrocytes (**Figure 5c**). The mDA specific genes that we examined in the ALN-neurons, including TH, DAT, DDC, VMAT2, PITX3, EN1 and GIRK2, were all increased in the presence of astrocytes Besides, genes involved in synaptic activities of the neurons, such as synapsin, synaptophysin, PSD95 and VAMP2, were also upregulated, indicating a promotion of astrocytes to the synaptic formation and function of the neurons [2, 33, 34]. On the other hand, not all genes examined for the SAB-astrocytes were improved by co-culturing with the ALN-neurons. A significant increase in gene expression was found for GFAP, GLT-1, AQP4 and GLUD2. It was found that the expression of GLAST in astrocytes was relatively constant whereas the expression of GLT-1 positively correlated with the maturation status [35, 36] and the synaptic activities of the neurons [37]. Therefore, an elevated expression of GLT-1 rather than GLAST, in line with an early study [26], indicated a more matured status of the SAB-astrocytes after co-culturing with neurons. Importantly, in the absence of direct contact, the increased maturation of both ALN-neurons and SAB-astrocytes were therefore evidenced by the presence of soluble factors secreted by the cells, known as the paracrine effect [33]. The transwell model is suitable for investigating neuroinflammation mediated by inflammatory cytokines. Evoked astrocytes by exposure of TNFα, IL-1α and C1q, which was categorized as A1 subtype, play a major role in CNS neuroinflammation [15]. We treated the SAB-astrocytes with TNFα and IL-1α and examined their gene profile and neural toxicity. It was shown that these astrocytes underwent a drastic morphological change and a slight increase in cell numbers after the stimulation (**Figure 6a**). QPCR also revealed a changed gene profile (**Figure 6b**). GFAP, S100β and AQP4 were significantly upregulated whereas GLT-1 and GDNF, representing the neural supportive role of the astrocytes, were significantly downregulated. Interestingly, for the comparison of astrocyte gene profile between the co-culture condition (**Figure 5c**) and the inflammatory condition (**Figure 6b**), several genes were constantly upregulated such as GFAP, S100β, AQP4 and GLUD2. We further introduced these inflammatory SAB-astrocytes to the ALN-neuron culture in the transwell model. Neuron death was observed 72 h after addition of these astrocytes, which was accompanied by greatly reduced complexity in the neural network formation (**Figure 6c**). Inflammatory astrocytes greatly undermined synaptic functions of the ALN-neurons, as was revealed by decreased gene expressions of synapsin, synaptophysin, PSD95 and VAMP2 (**Figure 6d**).

**Figure 6.**
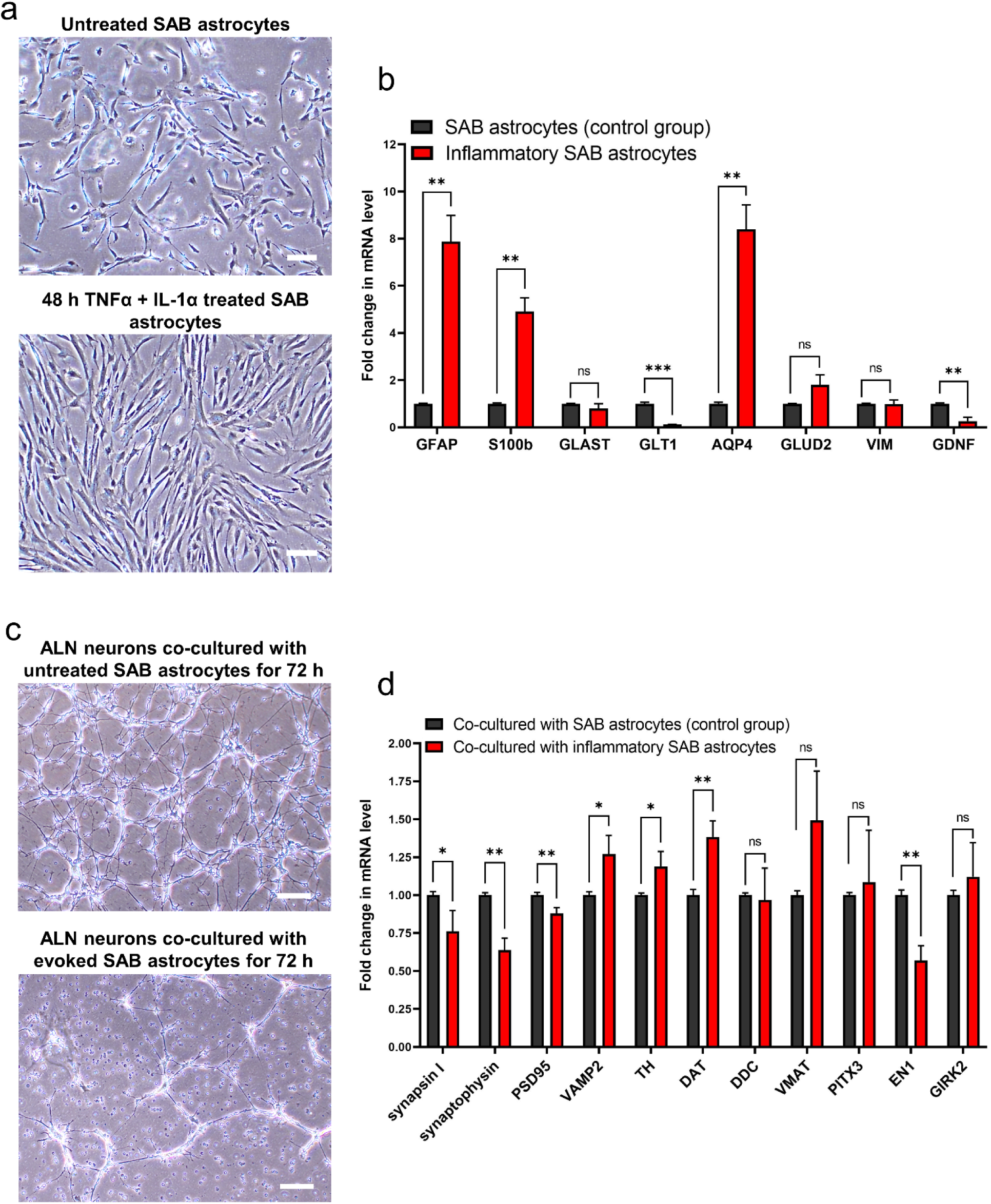
Transwell co-culture model consisting of ALN-neurons and inflammatory SAB-astrocytes. **(a)** Brightfield images showing the inflammatory SAB-astrocytes induced by TNFα and IL-1α. Scale bars are 200 µm. **(b)** QPCR quantification of the inflammatory SAB-astrocytes. **(c)** Brightfield images showing the ALN-neurons after co-culturing with inflammatory SAB-astrocytes in the transwell. Scale bars are 200 µm. **(d)** QPCR quantification of the ALN-neurons after co-culturing with inflammatory SAB-astrocytes. All graphs show mean ± SD. Statistical analysis was performed using student’s t-test; ns indicates no significant differences; \**P* < 0.05, \*\**P* < 0.01, \*\*\**P* < 0.001.

It has been found that the A1 astrocytes express a number of proinflammatory cytokines such as CXCL-, CCL-, IL-, C3a, GM-CSF [38–40], which compromise synaptic activities of the neurons as well as inducing neuron apoptosis [15, 39]. Given that the ALN-neurons were cultured without directly contacting the astrocytes in the transwell model, we confirmed that the death of the neurons was clearly caused by paracrine effect of the inflammatory astrocytes. Taken together, we showed the ALN-neurons and SAB-astrocytes were biologically responsive in the co-culture model. Besides, the SAB-astrocytes were functionally capable of inducing neuroinflammation.

## Discussion

The hiPSC-derived neural cells are valuable in both basic and translational studies. Importantly, neural cells derived from patient iPSCs represent an ideal platform for disease modeling and personalized medicine [20, 41–44]. Moreover, in the treatment of neurodegenerative diseases, autologous iPSCs are reliable cell sources for cell-based therapy. For example, as the midbrain nigrostriatal DA neurons are selectively dying in PD, cell-based therapy becomes the most promising and achievable treatment for this disease [45–50]. In this work, we explored the possibility to derive neurons and astrocytes from hiPSCs by two distinct methods. Even though essentially differing, both methods successfully generated the aimed cell types. In combination with other neural cell types as well as brain endothelial cells generated also from hiPSCs, these hiPSC-derived brain cells provide the opportunity for more authentic human BBB/NVU *in-vitro* modeling which favors both disease study and drug development [51].

### Dual-SMAD method

The induction of neuroectoderm fate is achieved through the inhibition of bone morphogenetic protein (BMP) and Activin/TGFβ signaling pathways, which is known as dual-SMAD inhibition [7, 17]. By simultaneously combining region-patterning morphogens, defined neural lineages can be obtained [17, 52, 53]. Although the dual-SMAD method has been well optimized and used extensively, it is considered rather laborious. Stepwise initiations of various expensive morphogens and growth factors are required during different time points of the differentiation. Besides, the concentrations for these factors should be precisely controlled. Moreover, these protocols normally take very long. For example, the prevailing mFPP-based mDA neuron differentiation protocols normally take 50 to 80 days [10, 18]. In our experiment, gene profiling showed that the mDA neurons from day 80 marked a more maturated stage than that from day 45, suggesting a long time should be waited before using these neurons as a qualified neural model. On the other hand, as gliogenesis has delayed onset than neurogenesis, the generation of astrocytes takes even longer time [54]. It was documented that the astrocyte differentiation protocols range mostly from 2 months to 6 months [17], with some of them even taking as long as one year [55]. Thus, the labor-intensive and time-consuming features of dual-SMAD protocols could greatly limit their use in certain applications.

### Direct lineage reprogramming

Direct lineage reprogramming is also known as transdifferentiation. By overexpressing neural lineage-restricted TFs, neurons or astrocytes can be rapidly generated from iPSCs [1, 41, 56, 57]. Strikingly, it was found that overexpression of a single TF, such as Ngn1/2 [31, 58] or ASCL-1 [30, 59, 60], was powerful enough to commit the neural fate. Meanwhile, by co-expressing other supporting TFs, purer and more matured neural subtypes can be then generated [30, 60, 61]. As activation of lineage TFs bypasses the neural progenitor/neural stem cell stage during the iPSC-to-neural conversion, such protocols could take very short time [62–64]. For example, neurons can be derived within a week [58, 60, 64, 65]. Notably, 7 days’ TF overexpression was found to suffice a stable neural conversion as endogenous TFs became prominent later on [34, 65].

On the other hand, TF overexpression even allows for direct neural conversion from somatic cells such as fibroblasts [38, 63–66]. Importantly, direct somatic-to-neural conversion is found to preserve age-associated epigenetics, which are important factors associated with many neurodegenerative diseases [41, 67–71]. Thus, for patient-specific disease modeling, this mechanism offers superior advantages over the use of patient’s iPSCs. In the current study, we overexpressed previously selected TF sets to derive the mDA neurons and astrocytes, respectively. In less than a month, both the differentiated ALN-neurons and SAB-astrocytes expressed their lineage-related markers. However, most of these reprogrammed neural cells were found to be rather immature. Differentiation of these cells still needs to be assisted with a co-culture strategy or with the addition of neurotrophic factors in the medium [30, 32, 60, 64, 72]. In line with it, we demonstrated a mutual maturation in both the ALN-neurons and SAB-astrocytes when cultured together. The co-culture model is not only important for neural maturation, but also it could reproduce complicated neural crosstalk in disease modeling. For example, co-culturing the neurons and astrocytes both generated from patient iPSCs has revealed complicated non-cell-autonomous effects that were involved in the disease pathogenesis [73]. To prove the usefulness of our co-culture model, we showed that it was convenient to model the neuroinflammation-induced neuronal degeneration. Notably, under inflammatory conditions, the astrocytes adopt an inflammatory phenotype whereas neurons are more prone to die. We noticed that the genes that have been upregulated in neurons when co-culturing with the supportive astrocytes were nonetheless deprived when co-culturing with the inflammatory astrocytes. These data indicated both the vulnerability of neurons and the double-faceted roles of astrocytes in affecting the neurons.

Despite many advantages of the direct reprogramming method in generating neural cells, there are also non-negligible drawbacks. The reprogramming efficiency appeared to be positively correlated with the expression level of the TFs [65]. Individual cells could have faced different dosing of each virus within the lentiviral cocktail, thus leading to variable expression of each TF which causes a high heterogeneity in the derived neural population. Furthermore, considering a single viral vector could successfully infect approximately 85% of the entire cell population, the optimal qual-infection percentage of a 4-viral combination (e.g., rtTA and 3 TFs) was estimated to be only 55% [13]. Above all, as overexpression of the lineage-restricted TFs bypasses the proliferative neural progenitor stage, it further decreases the yield of qualified neural cells. The low number of reprogrammed neural cells greatly hinders its application in large-scale disease modeling and drug testing experiments. In comparison, the dual-SMAD inhibition protocol, which allows the neural progenitors to be expanded at defined differentiation stages, could produce terminal neural cells for as many as 380-1780 times more than the original seeded number of the iPSCs [18, 50]. To solve these problems, multicistronic viral vectors can be used to increase the efficiency of equal TFs delivery [74]. Alternatively, stable iPSC clones that carry the DOX inducible TF-expressing cassettes can be generated [29, 65]. As these clones have been well selected beforehand, their differentiation efficiency and final yield of the neural cells were much higher compared to the iPSCs being directly infected [65].

Activation of endogenous TFs during *in-vivo* neurogenesis indeed follows a defined timeframe [75]. Unlike dual-SMAD method which active these TFs by stepwise giving related factors to the cells, almost lineage reprogramming protocols activate the TFs all at the beginning of differentiation. In terms of astrocyte generation, it was found that NFIA is necessary in driving early gliogenesis [34, 76, 77] whereas it is minimally required in matured astrocytes [35]. In line with this, several studies found that the prolonged overexpression of NFIA, other than NFIB or SOX9, prevented further maturation of those SAB-astrocytes [34, 38]. Nonetheless, this problem seemed to be tackled by applying only a short-term overexpression of these exogenous TFs, as this will awaken the endogenous ones which will be expressed during the rest of the differentiation in a more ontogenetic way.

Having the right regional identity in the derived neural cells is important for both disease modeling and cell-based therapy. It was found that astrocytes with regional specifications selectively support neurite growth and synaptic activity of the cultured neurons [78, 79]. Neurons with regional identities showed differentiated vulnerability to physiological and biochemical stresses [80, 81]. Importantly, implanted neurons with mismatched region specificity or subtypes failed to functionally integrate into the targeted area in the host brain [10, 52, 53, 82–84]. The dual-SMAD protocols were able to regionally pattern the derived neural cells, and these region identities could be preserved after *in-vivo* implantation [17, 27, 35, 50, 52, 85, 86]. Besides, the use of lineage-restricted iPSC lines allowed for more precise regional patterning [53]. Except for some cases [32], overexpression of a selected set of TFs was found to induce mixed region identities in the derived neural cells [30, 38, 87]. Interestingly, as researchers in the early days were dedicated in finding the TF combinations minimally required to generated functional neural cells from *in vitro*, some TFs that seemed unnecessary in promoting the neural differentiation but could be important in patterning the region specifications, may have been omitted. Even though the brain microenvironment could erase the wrong regional imprint of the implanted neural cells [88, 89], the mismatched regional marks in the *in-vitro* cultured cells could however lead to misleading data and conclusions. It is speculated the more TFs to be involved during direct reprogramming, the more explicit regional identity could be gained by the reprogrammed neural cells. Alternatively, TF-overexpression can be combined with other region patterning approach to derive region specificities [29, 90]. In principle, this hybrid method should also take the shortest differentiation time of the neural cells [90].

A nonnegligible problem of the lineage reprogramming methods might come from the use of lentiviruses. When transducing with lentiviruses, risks exist such as insertional mutations [91], transcriptome perturbation [92, 93] and tumorigenesis [94]. As a result, DNA integration-free delivery methods have been developed in recent years to generate either iPSCs or directly reprogrammed cells. These include non-integrating viral vectors [95, 96] or non-viral approaches [97] which include episomal DNA vectors [98], mRNA [99], miRNA [61], CRISPR/dCas9 [100], protein [62, 101] and small molecules [62, 102–106]. Such new approaches represent safer methods to derive the neural cells.

Today, one increasingly growing problem for the use of lineage reprogramming methods is the lack of standardized protocols [1, 56, 57, 107]. Notably, this is also true for the cytokine-based dual-SMAD methods [1, 19, 56, 107]. Lacking standardized protocols will result in unpredictable inconsistence in the phenotypes and biological behaviors of *in-vitro* neural models. Similarly, as a major component in the BBB, the generation of brain endothelial cells from hiPSCs also faces diverse protocols [108–110]. These are the main obstacles towards creating advanced and reliable *in-vitro* human neuronal/BBB/NVU models. Putting more effort into data comparison of these *in-vitro* derived neural cells between the lab and *in-vivo* will help to select the most faithful protocols and project their use towards biomedical applications.

## Conclusion

In this study, we differentiated the hiPSCs into functional mDA neurons and astrocytes. Two distinct differentiation methods were tested, and both showed successful derivation of the neurons and astrocytes. We were able to create an isogenic co-culture neural model by mixing these hiPSC-derived astrocytes and neurons. Importantly, the co-culture environment was proven necessary for the maturation of both cell types. The co-culture model also showed its convenience in investigating the astrocyte-induced neuroinflammation. These hiPSC-derived neural cells represented promising *in-vitro* platform for better dissecting human neurology.

## Author Approvals

All authors have seen and approved the manuscript, and that it hasn’t been accepted or published elsewhere.

## Acknowledgement

We acknowledge Ruben de Vries and Sibel Bahtiri, who were master students enrolled in the Drug Innovation program at Utrecht Institute for Pharmaceutical Sciences (UIPS), for providing experimental assists in this work.

## Supplementary information

### Supplementary methods

#### Cell culture

Mouse embryonic fibroblasts (MEFs) were derived from E14.5 mouse embryos. MEF medium consists of high glucose DMEM (#10313021, Thermo Fisher) supplemented with 10% FBS, 0.1 mM NEAA, 1 mM sodium pyruvate and 1 × Antibiotic Antimycotic. Upon reaching confluency of 80-90%, cells were detached with 0.025% trypsin EDTA and sub-cultured at the ratio of 1:5 in new flasks.

Human primary somatic fibroblasts were isolated from healthy donors and were maintained in culture flasks in 5% CO_2_ humidified incubator at 37 ℃. Human fibroblast medium consists of DMEM supplemented with 10% FBS and 1 × Antibiotic Antimycotic. Upon reaching confluency of 70-80%, cells were detached with 0.025% trypsin EDTA and sub-cultured at the ratio of 1:10 in new flasks.

### Antibiotic kill curve on hiPSCs

Single hiPSCs were prepared as described earlier. A total of 10,000 cells were seeded per well in Matrigel coated 96-well plates. Next day, medium was switched to either the neuron induction medium or astrocyte induction medium, supplemented with 10 µM Y27632. The day after, puromycin (P9620, Sigma-Aldrich) prepared at serial concentrations were given to the cells in fresh medium without Y27632. During the selection period, antibiotic and cell medium was refreshed daily. On indicated days, the alamarblue assay was performed to determine cell viability (**Figure S1a**). Briefly, cells were incubated with cell medium containing 50 μM Resazurin (R7017, Sigma-Aldrich) for 4 h. The medium was then collected and centrifuged. The supernatant was transferred to a fluorescence assay plate and the fluorescence intensity was measured at 530/590 nm (excitation/emission) using a fluorescence plate reader (Jasco). The antibiotic kill curve was plotted using GraphPad software. The lowest puromycin concentration that killed 95-100% cell population on the desired day was selected for the hiPSC differentiation experiment.

---

**Figure S1.**
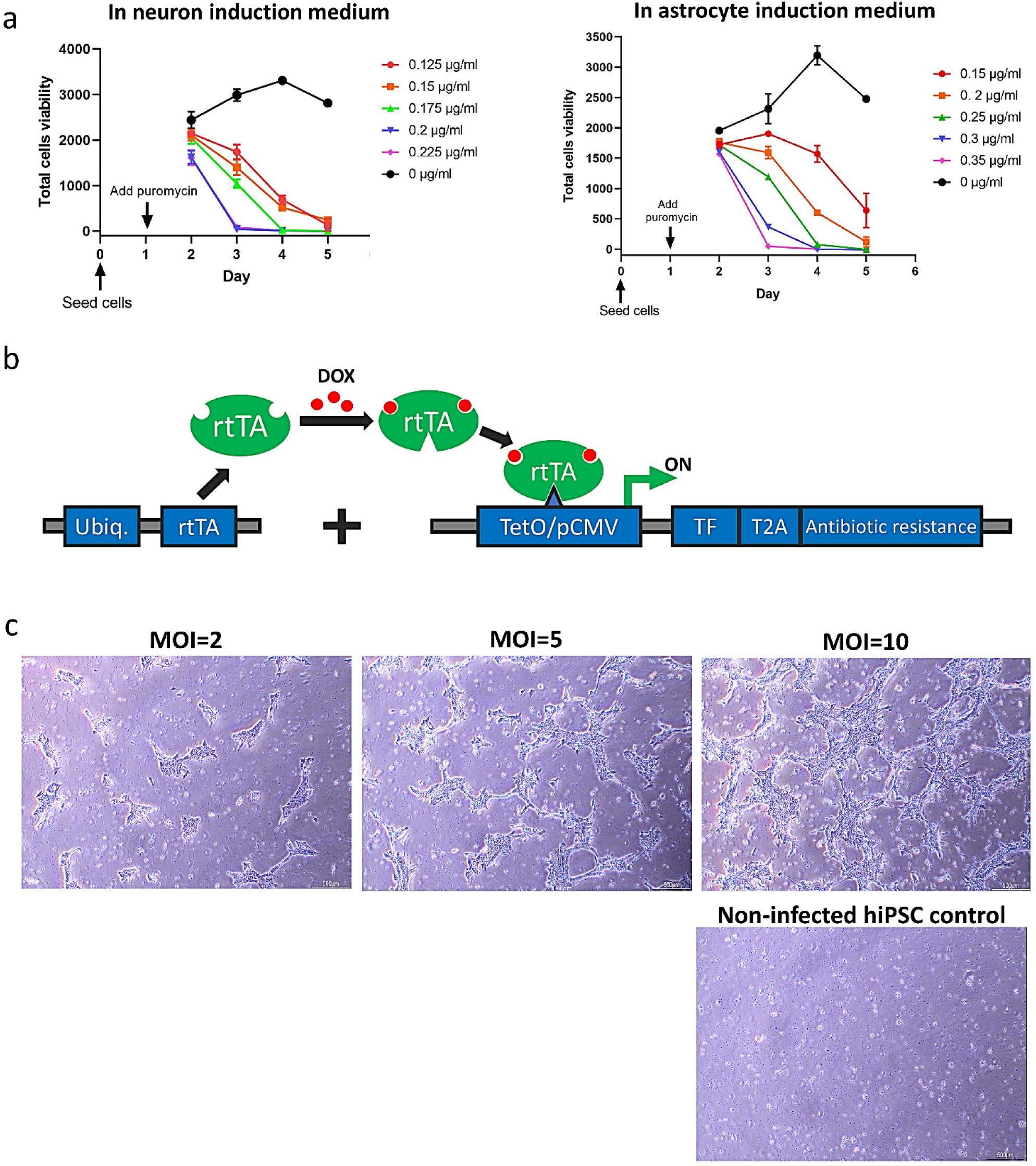
Antibiotic kill curve on hiPSCs. **(a)** Puromycin selection on hiPSCs cultured in neuron induction medium (left) or astrocyte induction medium (right). **(b)** Schematic illustration of the tet-On expression system. The system contains at least two vectors. The reverse tetracycline-controlled trans-activator (rtTA) is expressed under the control of ubiquitous promoter. Gene of interest is expressed under the control of tet-on operator (tetO). In the presence of DOX, the rtTA-DOX complex binds to the tetO operator and activates gene expression. **(c)** Brightfield images showing the end of a 4-day puromycin selection on hiPSCs infected with ALN and rtTA viruses under different MOI.

**Figure S2.**
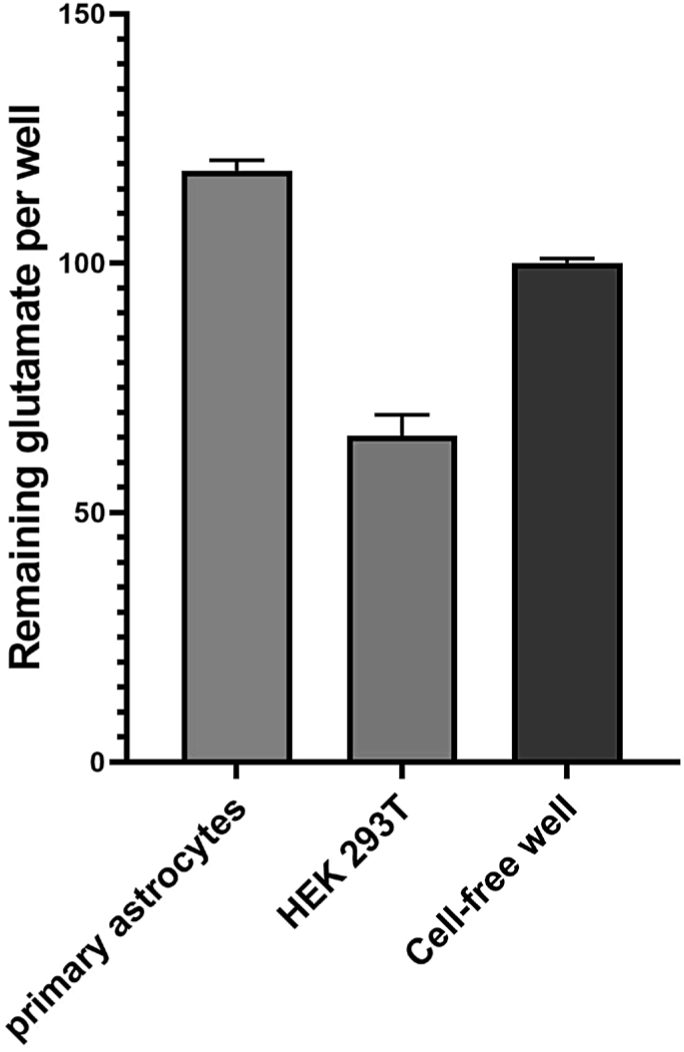
Glutamate uptake of primary human astrocytes. HEK 293T cells were used as control cells. Cell-free wells were used as blank. Graph shows mean ± SD.

**Table S1.** Lentiviral plasmids used in this study.

| Plasmid | Addgene catalog number |
| --- | --- |
| pMDLg | #12251 |
| pRSV-Rev | #12253 |
| pMD2.G | #12259 |
| FUW-M2rtTA | #20342 |
| TetO-FUW-NFIA (Nuclear Factor I A) | in-house |
| TetO-FUW-NFIB (Nuclear Factor I B) | in-house |
| TetO-FUW-SOX9-puro (SRY-Box Transcription Factor 9) | #117269 |
| TetO-FUW-ASCL1-puro (Achaete-Scute Family BHLH Transcription Factor 1) | # 97329 |
| TetO-FUW-NURR1 (Nuclear receptor-related 1 protein) | in-house |
| TetO-FUW-LMX1A (LIM Homeobox Transcription Factor 1 A) | in-house |

**Table S2.**
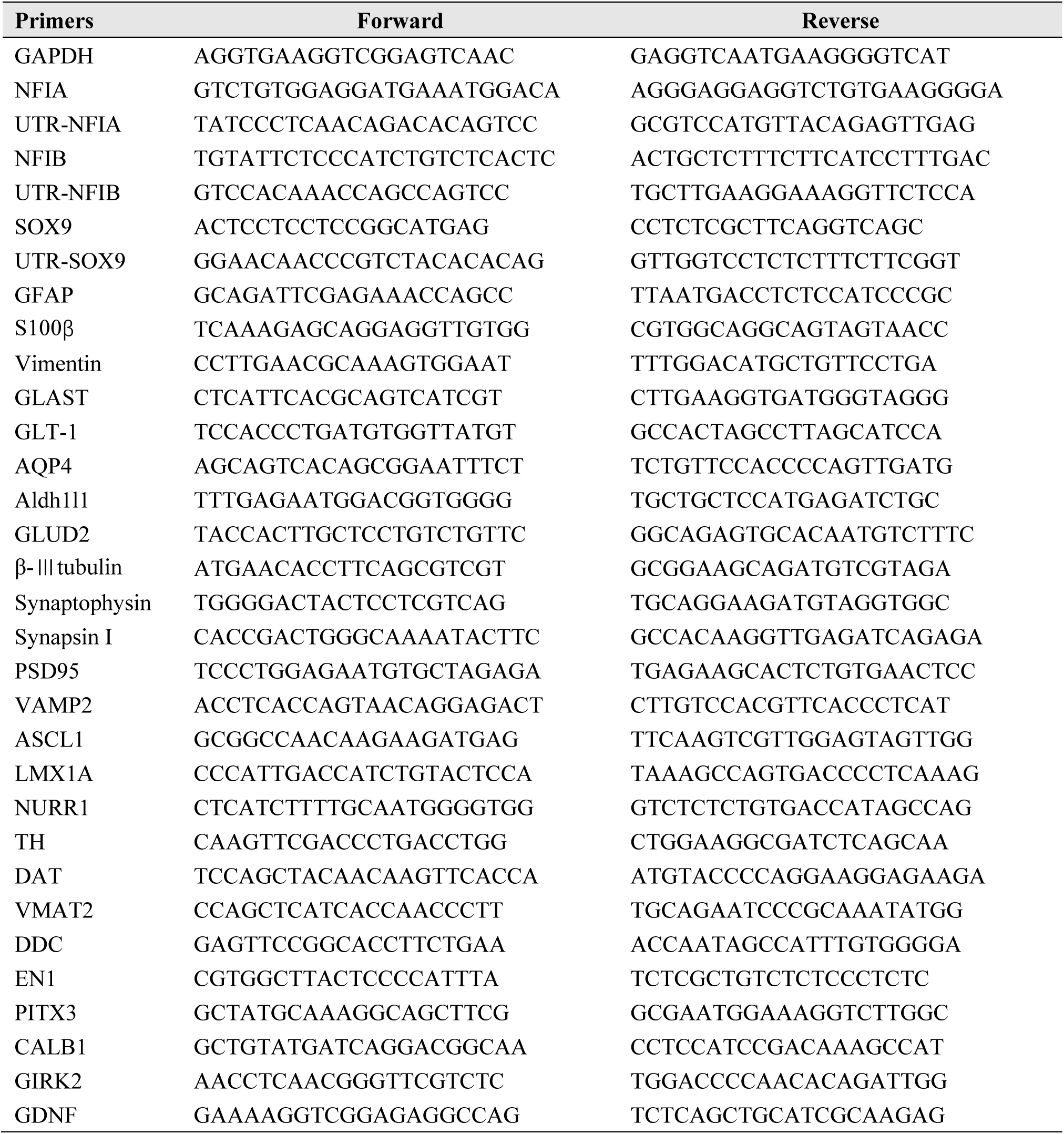
The qPCR primers used in this work.

